# Optimization and quantitative evaluation of fluorescence *in situ* hybridization chain reaction in clarified fresh-frozen brain tissues

**DOI:** 10.1101/763920

**Authors:** Vivek Kumar, David M Krolewski, Elaine K Hebda-Bauer, Aram Parsegian, Brian Martin, Mathew Foltz, Huda Akil, Stanley Watson

## Abstract

Transcript labelling in intact tissues using *in situ* hybridization chain reaction has potential to provide vital spatiotemporal information for molecular characterization of heterogeneous neuronal populations. However, it remains relatively unexplored in fresh-frozen brain which provides more flexible utilization than perfused tissue. In the present study, we optimized the combination of *in situ* hybridization chain reaction in fresh-frozen rodent brains and then evaluated the uniformity of neuronal labelling between two clearing methods, CLARITY and iDISCO^+^. We found that CLARITY yielded higher signal-to-noise ratios but more limited imaging depth and required longer clearing times, whereas, iDISCO^+^ resulted in better tissue clearing, greater imaging depth and a more uniform labelling of larger samples. Based on these results, we used iDISCO^+^-cleared fresh-frozen rodent brains to further validate this combination and map the expression of a few genes of interest pertaining to mood disorders. We then examined the potential of *in situ* hybridization chain reaction to label transcripts in cleared postmortem human brain tissues. The combination failed to produce adequate mRNA labelling in postmortem human cortical slices but produced visually adequate labeling in the cerebellum tissues. We next, investigated the multiplexing ability of *in situ* hybridization chain reaction in cleared tissues which revealed inconsistent fluorescence output depending upon the fluorophore conjugated to the hairpins. Finally, we applied our optimized protocol to assess the effect of glucocorticoid receptor overexpression on basal somatostatin expression in the mouse cortex. The constitutive glucocorticoid receptor overexpression resulted in lower number density of somatostatin-expressing neurons compared to wild type. Overall, the combination of *in situ* hybridization chain reaction with clearing methods especially iDISCO^+^ may find broad application in the transcript analysis in rodent studies but its limited use in postmortem human tissues can be improved by further optimizations.

## Introduction

Ribonucleic acids, especially mRNA and non-coding RNAs, give cells their molecular identity through their unique combination and can serve as important markers of activity-dependent gene expression and disease conditions. Since its first introduction, *in situ* hybridization has been a valuable tool to study RNA biology (Pardue and Gall 1969). Fluorescence *in situ* hybridization (FISH), in particular, has potential to provide high resolution spatiotemporal information for the molecular characterization of heterogeneous cell populations in the brain. Notably, recent advances in RNA/DNA hybridization techniques allow mapping of multiple mRNA targets simultaneously and with subcellular-specificity at both organ level (e.g. brain, kidney, liver) as well as organism (e.g., zebrafish) level. One such approach is the ‘*in situ* hybridization chain reaction’ based FISH (HCR FISH) (Dirks and Pierce 2004; Choi et al. 2010; Choi et al. 2018). In principle, self-assembly of metastable fluorophore-labeled DNA hairpins into tethered fluorescent amplification polymers through a chain reaction, is triggered by the presence of an initiator strand (Dirks and Pierce 2004). Applying several of these custom probes to tissue concurrently allows for multiplexing of up to five targets with high specificity. Moreover, the relatively short lengths of probes and hairpins make this technique particularly favorable for intact tissue transcriptional analysis.

The advent of the HCR FISH method alongside the recent development of intact tissue clearing and high-throughput microscopic imaging methods has broadened the path to obtain unprecedented information about the molecular identity, interactions, and activity-dependent functions in the brain. To date, several tissue clearing approaches have been used to achieve rodent brain transparency but only a few have proved to be versatile and successful across a wide range of applications (Silvestri et al. 2016; Azaripour et al. 2016; Wan et al. 2018). Based on factors like clearing capability, target biomolecule fixation, ability to perform ex-vivo labeling, fluorescence retention and imaging depth, we chose to explore two methods 1) tissue lipid matrix replacement-based CLARITY (Chung et al. 2013) with modifications (Tomer et al. 2014; Yang et al. 2014; Zheng and Rinaman 2016; Krolewski et al. 2018) and 2) non-aqueous solvent-based iDISCO^+^ (Renier et al. 2014). There has been pioneering work done (Sylwestrak et al. 2016; Kramer et al. 2018; Park et al. 2018) but the combination of HCR FISH with clearing techniques still remains relatively unexplored especially with fresh-frozen tissues. Flash-freezing fresh rodent brains provides flexible utilization for studying a wide array of biomolecules (e.g., RNA, protein, metabolite) using multiple methods like *in situ* hybridization, immunohistochemistry, PCR, western blotting, spectroscopy. Traditionally, fresh-frozen tissues have been preferentially used as starting material for both radioactive- and digoxigenin-based *in situ* hybridization. One of our major research interest involves postmortem human brain which are certainly a valuable source of information for psychiatric and neurodegenerative disorders. These tissues are routinely collected without any perfusion fixation and then stored frozen. Based on the unique opportunity fresh-frozen tissues provide, we decided to investigate the compatibility and efficiency of HCR FISH with two clearing approaches, CLARITY and iDISCO^+^ in fresh-frozen tissues. Thus, the following experiments were designed to 1) optimize the combination of HCR FISH with CLARITY and iDISCO^+^ in fresh-frozen rodent brains and quantify the somatostatin (*Sst*) FISH signal for the uniformity of cellular labelling between the two clearing methods, 2) use HCR FISH to validate and map the expression of a few genes of interest in fresh-frozen rodent and postmortem human brain tissues, 3) investigate the efficiency of HCR FISH to detect multiple genes in cleared samples, and 4) apply an optimized volumetric quantitation pipeline of HCR FISH in cleared tissues to compare the basal level expression of the *Sst* mRNA in the cortex of wild type (WT) and glucocorticoid receptor over-expressing (GRov) mice (Wei et al. 2004; Wei et al. 2007; Hebda-Bauer et al. 2010; Wei et al. 2012).

## Experimental Procedures

### Animal care

All animal care and experimental procedures followed the guide for the Care and Use of Laboratory Animals: Eighth Edition (revised in 2011, published by the National Academy of Sciences), and approved by the University of Michigan Committee on the Use and Care of Animals. The generation of GRov mice has been described previously (Wei et al. 2004). The GRov mouse line was established in the Akil laboratory by breeding founders and their progeny to C57BL/6J mice, and all transgenic mice are maintained as hemizygotes. Adult male Sprague-Dawley rats (3-4 months old) were obtained from Charles River Laboratories (Wilmington, MA) and acclimated to their cages for one week. Mice and rats were housed on a 14:10 and 12:12 light/dark cycle, respectively, with ad libitum access to food and water. Adult male GRov and WT littermates (2-4.5 months old) were sacrificed by rapid decapitation, and their brains removed, snap frozen in 2-methyl butane (Isopentane, cooled with dry ice), and stored at −80°C prior to use.

### Human brain tissue acquisition

Blocks of temporal cortex and cerebellum from postmortem human brains were used, which were collected by the Brain Donor Program at the University of California, Irvine with the consent of the relatives of the deceased. Brains were removed at autopsy, quickly chilled to approximately 4°C and then cut into series of 0.75 cm thick coronal slices that were quickly frozen and stored at −80°C as previously described (Jones et al. 1992). Slabs were then placed on dry ice blocks and dissected using a fine-toothed saw to generate tissue blocks of approximately 4×3 cm in size that were then stored at −80°C until further use. Information regarding physical health, medication use, psychopathology, substance use, and details about the final hours of the deceased was obtained from medical records, the coroner’s investigation, the medical examiner’s conclusions and interviews with relatives. Brain blocks from the subjects with a diagnosis were used in this study. All four male subjects were between 35-66 yrs. of age, and their blood pH during the brain collection ranged from 6.8 to 7.2.

### Tissue preparation

#### Immersion-fixation in paraformaldehyde (PFA)

Fresh-frozen rodent brains and postmortem human brain blocks were immersion-fixed in 4% PFA at 4°C for 12-36 h depending upon thickness.

#### CLARITY sample preparation

Tissues were prepared as described in (Tomer et al. 2014; Krolewski et al. 2018) with modifications to immersion-fixation and the FISH protocol. After fixation, tissues were rinsed in boric acid buffer (0.2 M borate/0.1% Tween-20, pH 8.5) and then immersed in 2% hydrogel solution (2% acrylamide, 0.25% VA-044 initiator, 1x phosphate buffer-saline (PBS), 4% PFA) for 24 h at 4°C. Tubes containing tissues were vacuum degassed then incubated for hydrogel polymerization at 37°C for 3-5 h. Fixed tissues were washed in boric acid buffer for 3 × 2 h each then sectioned using a vibratome (500-1200 µm thick) or a rodent brain matrix (1-4 mm thick). Cut slices were then passively cleared in 4% sodium dodecyl sulfate (SDS) at 45°C for 1-3 weeks and rinsed in boric acid buffer with 0.1 % tween-20 for 3 × 2 h before processing for FISH.

#### iDISCO^+^ sample preparation

Tissues were prepared as described in (Renier et al. 2014) with modifications for immersion-fixation and the FISH protocol. Fixed samples were washed in 0.01M PBS/0.1% tween-20 (PBSTw) for 3 × 1 h, with an additional overnight wash at room temperature. Tissues were then sectioned using a vibratome (500 µm-1 mm) or a brain matrix (1-4 mm). Additionally, an intact hippocampus from both hemispheres were dissected from a rat brain after the PFA fixation and rinsing. Tissues were then dehydrated in 20%, 40%, 60%, and 80% methanol (in ddH_2_O) for 1-2 h each, and 100% methanol for 2 × 1-3 h each. Samples were bleached with 5% H_2_O_2_ in ice cold methanol at 4°C overnight, and then rehydrated in 80% methanol/H_2_O for 1 h, 50% methanol/H_2_O for 1 h, and finally in PBSTw for 1 h twice. Samples were then used for HCR FISH.

### Probe design and synthesis

Split-initiator DNA probes were either purchased from Molecular Instruments, Inc. (Los Angeles, California, USA) or designed in our lab based on Choi et al. 2014, 2018 and synthesized by Integrated DNA Technologies (Coralville, Iowa, USA). In this study, probes for the following genes were used 1) Rat: *somatostatin* (*Sst*, NM_012659.2), *parvalbumin* (*Pvalb*, NM_022499.2), *dopamine-beta-hydroxylase* (*Dbh*, NM_013158.2), *tyrosine hydroxylase (Th*, NM_012740.3), *solute carrier family 6 member 3 (dopamine transporter, Slc6a3*, NM_012694.2); 2) Mouse: somatostatin (*Sst*, NM_009215.1); 3) Human: *somatostatin* (*SST*, NM_001048.4), *parvalbumin* (*PVALB*, NM_002854.2) and *calbindin-1* (*CALB*, NM_001366795.1). DNA hairpins conjugated with AlexaFluor-488 (AF-488), AF-594 and AF-647 were purchased from Molecular Instruments, Inc. (Los Angeles, California, USA). Additionally, DNA hairpins conjugated with fluorescein isothiocynate (FITC), cyanine-3 (Cy3), and AF-647 were designed based on Choi et al. 2014 and synthesized and conjugated by Integrated DNA Technologies (IDT).

### HCR FISH

We modified the HCR FISH method as described by Choi et al. 2014, 2018 for thick fresh-frozen samples. CLARITY- and iDISCO^+^-processed samples were equilibrated with 5x sodium citrate/0.01% tween-20 (SSCTw) buffer for 1 h, then acetylated with 0.25% v/v acetic anhydride solution for 30-60 min. After briefly rinsing with ddH2O, samples were then equilibrated in hybridization buffer (30% deionized formamide, 5x SSC, 0.5 mg/ml yeast tRNA, 10% dextran sulfate) for 1 h. Tissues were then incubated in the hybridization buffer containing 1-6 nM initiator-labeled probes at 37°C for 12-72 h depending upon the tissue thickness. Following hybridization, samples were washed at 37°C with probe wash buffer (30% formamide, 5x SSC, 9 mM citric acid, 0.1% tween 20) for 3 times and then twice with 5x SSCTw for 1 h each. For samples thicker than 1 mm, an additional rinse of 12 h in 5x SSCTw at room temperature was also performed. Tissues were equilibrated in amplification buffer for 1-3 h (5x SSC, 10% dextran sulfate, and 0.1% Tween 20). Fluorophore-labeled hairpins were diluted separately from 3 µM stock to a 2.25 µM final concentration in 20x SSC, heated at 90°C for 90 seconds, and then cooled to room temperature for 30 min in the dark. Cooled hairpins were diluted to a 20-60 nM final concentration in amplification buffer. Tissues were incubated in amplification buffer with hairpins for 1-3 days at room temperature. Finally, tissues were washed in 5x SSCTw for 3 × 1 h (overnight 1 × wash for sections over 1 mm). For *Sst* quantitation studies, we used a final concentration of 2 nM probe and 20 nM hairpins based on concentration gradient study to achieve a median saturation of the *Sst* signal.

### Refractive index (RI) matching

After FISH, CLARITY samples were optically matched by incubating in 88% histodenz (Sigma-Aldrich, St. Louis, Missouri, USA) at 37°C overnight (Yang et al. 2014). iDISCO^+^ samples were dehydrated in 20%, 40%, 60%, 80%, and 100% methanol (in ddH_2_O) for 1 h each with an additional 1-2 h or overnight dehydration in 100% methanol. Samples were then incubated in 1 volume of methanol and 2 volumes of dichloromethane (DCM, Sigma-Aldrich) until they sank in glass vials. Residual methanol from tissues was then washed by two changes of 100% DCM for 30 min. each. Finally, samples were incubated (without shaking) in dibenzyl ether (DBE, Sigma-Aldrich) for 30-60 min to match the RI.

### Confocal Microscopy- Imaging and visualization

Samples were placed in 88% histodenz- or DBE-filled chambers made with 3M™ mounting putty on a glass slide and cover-slipped. Image stacks were acquired on an Olympus Fluoview 1000 confocal microscope using a 10x/0.4 N.A. objective (WD: 2.2 mm) at 405 nm (DAPI), 488 nm, 594 nm, and 647 nm excitation. Image dimension are 1024 × 1024 (1.24 µm/pixel × 1.24 µm/pixel) in the xy-plane with *z*-step of 4.26 µm. For comparative quantitation of mouse *Sst* signal, images were acquired within a region of interest (ROI) consisting of cingulate and primary/secondary motor cortex (rostral) to retrosplenial agranular and retrosplenial granular cortex (caudal) and the ROI was kept consistent for all the samples. The image acquisition settings (i.e., voltage (HV, value=720), gain (value=0), and offset (value=5)) in the Fluoviewer software of the confocal microscope were kept constant for quantitative comparisons. Three-dimensional volume rendering and ortho-slice/*xy*-pane mode visualization was performed using Amira (ThermoFisher Scientific, Waltham, Massachusetts, USA) and Imaris (Bitplane Inc., Concord, Massachusetts, USA). Olympus FluoViewer software was used to extract histogram data for the confocal microscope-acquired images.

### Light sheet microscopy- Imaging and visualization

Imaging of large intact samples was performed on the CLARITY-optimized light-sheet microscope (COLM) (Tomer et al. 2014; Krolewski et al. 2018). RI-matched samples were mounted in a quartz cuvette and filled with 88% histodenz or DBE. The entire coronal slice was acquired in a series of tiles with a 10x/0.6-NA objective (WD: 3 mm). Voxel dimension of the image stacks are 0.595 µm/pixel × 0.595 µm/pixel in *xy*-plane and 5 µm/slice in the *z*-direction. The number of tiles was specified by defining the coordinates of the two opposing corners of the entire tissue slice plus a 15-20% tile overlap. Following the optimization of light-sheet and focal-plane alignment parameters over the entire sample space, final image acquisition was initiated and data collected. Tiles of acquired image stacks were stitched together using an in-house modified version of Terastitcher (Bria and Iannello 2012). Stitched coronal image volumes were 3D rendered and visualized using Amira and Imaris. ImageJ was used to extract histogram data from the COLM-acquired images.

### Quantitation and statistical analysis

Quantitative analysis of *Sst* transcript expression in the mouse cortex was performed on the confocal microscope-acquired image stacks using Amira software. Imaging was systematically randomized among the samples to minimize the variation resulting from longer incubation in the histodenz or DBE. The analysis pipeline involved noise removal, thresholding, and quantitation. Image stacks were first processed to achieve optimal thresholding by applying a median filter to both CLARITY and iDISCO^+^ images. Individual *xy*-planes of both CLARITY and iDISCO^+^ stacks were median filtered to reduce the contrast and soften the edges of objects. Although this process reduces the contrast and tends to defocus the image, it helps to eliminate the background noise which substantially improves the thresholding. Processed images were then locally segmented through each *xy*-plane to extract the target neurons. The extent of thresholding was proportionally determined based on the mean tissue background intensity and validated manually across z-planes and samples to make sure that the majority of the segmented objects are neuronal cell bodies. Further, in order to include only intact cells and not ‘poorly’ fragmented objects, the output was filtered by a combination of morphometric parameters: area, gray level intensity, perimeter, length, breadth, width, and circularity. These morphometric cutoff values were manually determined and were different for CLARITY and iDISCO^+^ samples due to tissue expansion and shrinkage, respectively, but kept consistent across all the samples in each clearing group. Finally, independent sample t-tests (to compare the means between 2 groups) or one-way ANOVAs with Tukey-alpha post hoc tests (to compare the means between multiple groups) were performed. Data are represented as mean ± SD.

## Results

### Optimization and comparison of HCR FISH signal between CLARITY and iDISCO^+^

The HCR FISH method as described by Choi et al. 2014, 2018, was modified to include an incubation step of acetylation with acetic anhydride solution which improved the signal to noise (S/N) ratio (data not shown). For iDISCO^+^ processing, we omitted the tissue permeabilization step with dimethylsulfoxide (DMSO)/glycine/Triton x-100 as mentioned in Renier et al. 2014. This step did not improve the probe permeability, but instead appeared to adversely affect the hybridization step which could be related to the ability of DMSO to form hydrogen bonds with nucleobases and promote destacking, especially in single stranded RNAs (Lee et al. 2013). Following the successful optimization, we compared the efficiency of HCR FISH between CLARITY (n=4) and iDISCO^+^ (n=5) samples by analyzing the *Sst* mRNA labelling in the cortex of WT mice (Fig. 1a *vs.* 1b; supplemental movie ESM_1). We found that the mean intensity of detected *Sst*^+^ neurons is similar between the CLARITY and iDISCO^+^ samples, with relatively higher variation in the iDISCO^+^ group (Fig. 1c). However, the mean tissue intensity is significantly lower in CLARITY samples *vs.* iDISCO^+^ (Fig. 1d) when calculated as background intensity with *Sst* signal (two-tailed t(7) = −4.08, p = 0.005) or without (two-tailed t(7) = −5.33, p = 0.001). Within a consistently acquired ROI volume, the CLARITY image stacks comprised a significantly lower number of *Sst*^+^ neurons per unit volume (two-tailed t(6) = −8.32, p = 0.000; Fig. 1e) *vs.* iDISCO^+^. In addition, the detected *Sst*^+^ neurons in CLARITY samples were comprised of significantly higher mean number of pixels *vs.* iDISCO^+^ (two-tailed t(7) = 13.16, p = 0.000) (Fig. 1f). Altogether, these measurements reproduced and validated the previously known expansion in CLARITY samples and shrinkage caused by iDISCO^+^ processing, while HCR FISH intensity for *Sst* labeling remains comparable between the two methods.

**Fig. 1.**
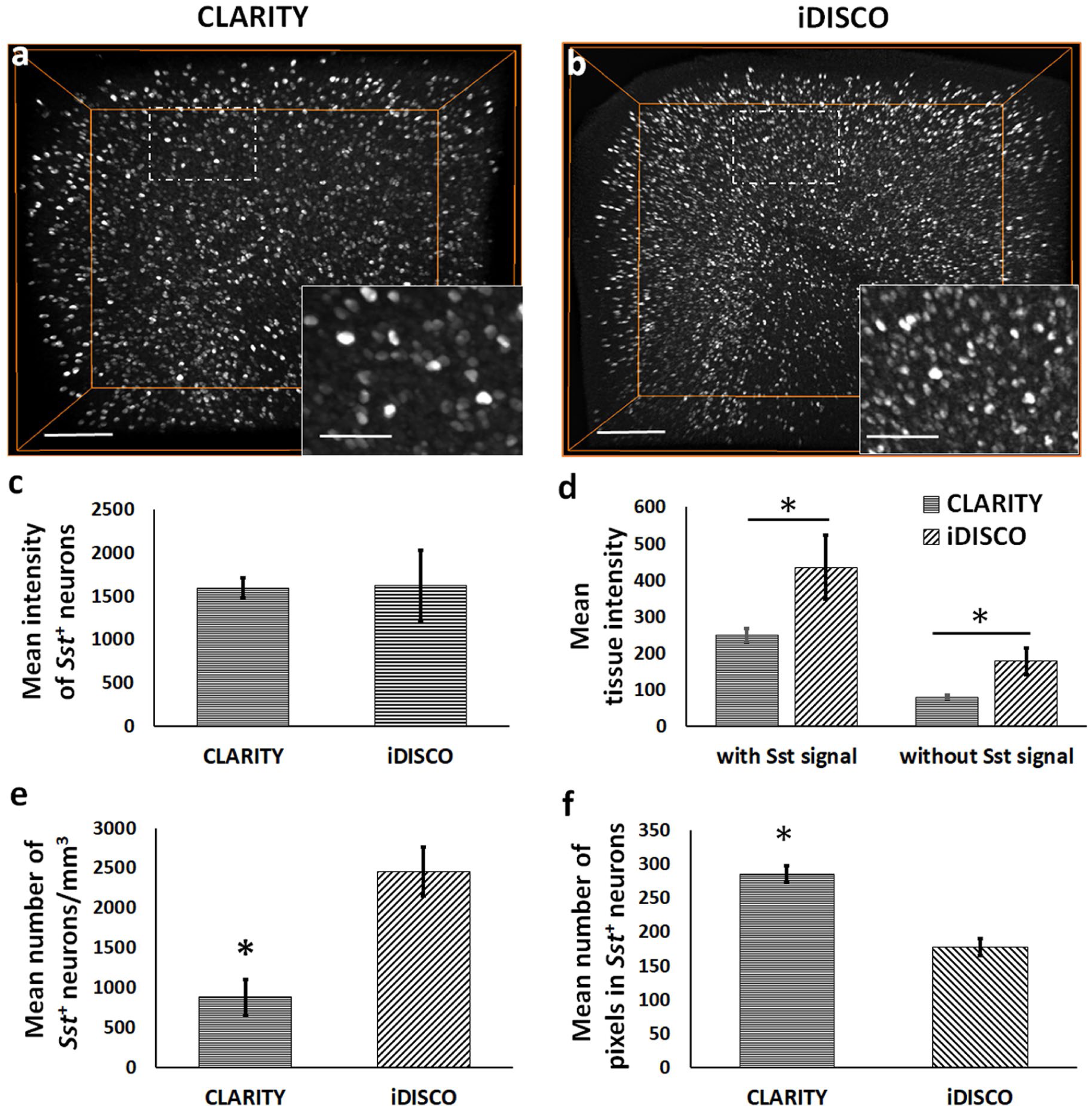
Quantitative differences in the HCR FISH signal (*Sst* transcript expression) between CLARITY and iDISCO^+^. (a-b) 3D volume-rendered images of representative CLARITY and iDISCO^+^ samples, respectively, showing the observed differences in the detected neuronal density and mean tissue intensity. (c) Bar-diagram shows comparable mean intensity of the detected *Sst*^+^ neurons in the acquired cortical ROI between CLARITY and iDISCO^+^. In CLARITY samples, mean background tissue intensity, with or without including the *Sst* signal, and (e) mean number of *Sst*^+^ neurons per unit volume were significantly lower compared to the iDISCO^+^ samples, for consistent cortical ROIs. (f) Bar-diagram shows a significantly higher mean number of pixels in the detected *Sst*^+^ neurons in CLARITY images *vs.* iDISCO^+^. *p< 0.05. Scale bars-200 µm (Insets-75 µm).

We next compared the fluorescence signal as a function of *z*-depth to assess any difference which could result from the combination of tissue permeability for DNA probes and optical transparency for imaging between CLARITY and iDISCO^+^ samples. Confocal microscope-acquired image stacks revealed an attenuation of signal in CLARITY samples with increasing depth (Figs. 2a; 2c *vs.* 2d; left panel in supplemental movie ESM_1), whereas iDISCO^+^ samples had a relatively uniform labelling across the z-depth (Figs. 2b; 2e *vs.* 2f; right panel in supplemental movie ESM_1). In CLARITY samples, thresholded neurons in deeper layers consist of a significantly lower number of pixels in comparison to those in upper layers (F (5, 47) = 17.806; 100 µm *vs.* 500 µm: p = 0.008, 100 µm *vs.* 600 µm: p = 0.001; Figs. 2g *vs.* 2h; 2k). In contrast, the mean number of pixels remained relatively consistent among the thresholded top and bottom layer neurons in the iDISCO^+^ samples (Figs. 2i *vs.* 2j; 2k). In summary, the decrease in the optical accessibility and, in turn, the reduced pixel intensity with ***z***-depth potentially result in the detection of fewer numbers of neurons in CLARITY compared to iDISCO^+^ samples.

**Fig. 2.**
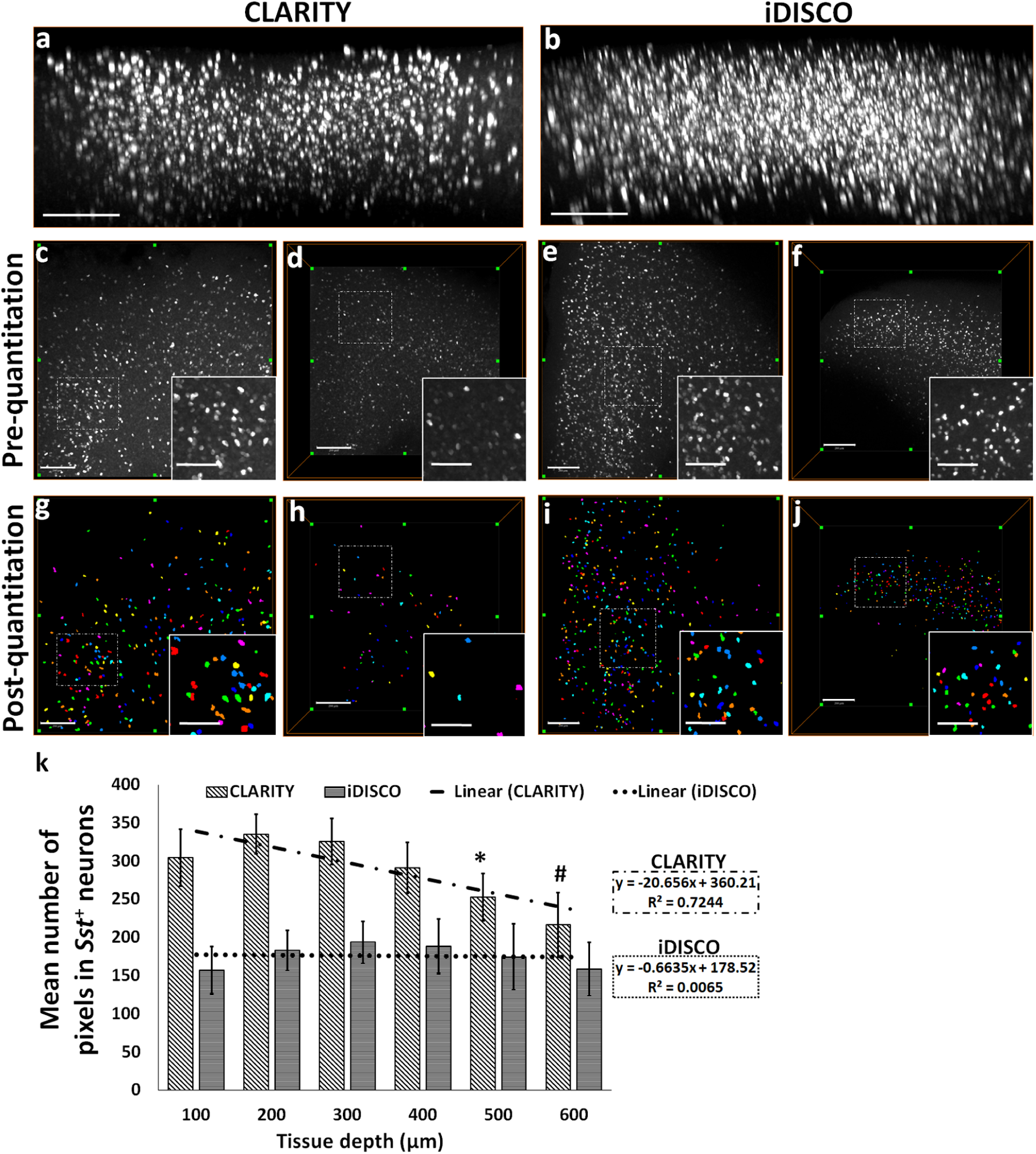
Attenuation of the HCR FISH signal in CLARITY samples measured as a function of z-depth. *xz*-plane view of 3D rendered confocal microscope-acquired volumes of (a) CLARITY and (b) iDISCO^+^ showing greater attenuation of HCR FISH signal with depth. Representative ***xy***-plane images at 100 µm z-depth: CLARITY-(c) pre-quantitation, (g) post-quantitation, iDISCO^+^-pre-quantitation, (i) post-quantitation, respectively. Representative ***xy***-plane images at 500 µm z-depth: CLARITY-(d) pre-quantitation, (h) post-quantitation, iDISCO^+^-(f) pre-quantitation, (j) post-quantitation, respectively. (k) Bar-diagram shows mean number of pixels in the detected neurons as a function of increasing *z*-depth (100 to 600 µm) in CLARITY and iDISCO^+^ samples. For CLARITY samples, *p<0.05 for comparisons between 100/200/300 *vs.* 500 µm and #p<0.05 for 100/200/300/400 *vs.* 600 µm. Scale bar-(a-b) 300 µm; (c-j) 250 µm (Inset bars-100 µm).

### HCR FISH in large iDISCO^+^-cleared fresh-frozen rodent and postmortem human brain

High resolution volumes acquired on COLM allowed us to visualize and reproduce the previously reported anatomical distribution of a few genes of interest to our lab. Layer specific expression of *Sst* in cortical slices of rat (Fig. 3a, supplemental movie ESM_2) shows a relatively higher density of *Sst* interneurons (Martinotti cells) in the superficial (L2/3) and deep (L5/6) cortical layers. Midlayer (L4) contains a sparse number of *Sst*^*+*^ neurons mainly corresponding to small basket cells (Iritani and Satoh 1991; Kawaguchi and Kubota 1997; Markram et al. 2004; Urban-Ciecko and Barth 2016). Fig. 3b and supplemental movie ESM_3 show the *Sst* mRNA expression in a COLM-acquired volume of an intact rat hippocampus which was dissected from the brain post-PFA fixation and then processed through iDISCO^+^ and HCR FISH. Fig. 3c and supplemental movie ESM_4 show the *Sst* mRNA expression in a representative COLM-acquired volume of a whole mouse hemisphere, with supplemental movie ESM_5 showing a zoomed-in view of the expression in the brainstem region.

**Fig. 3.**
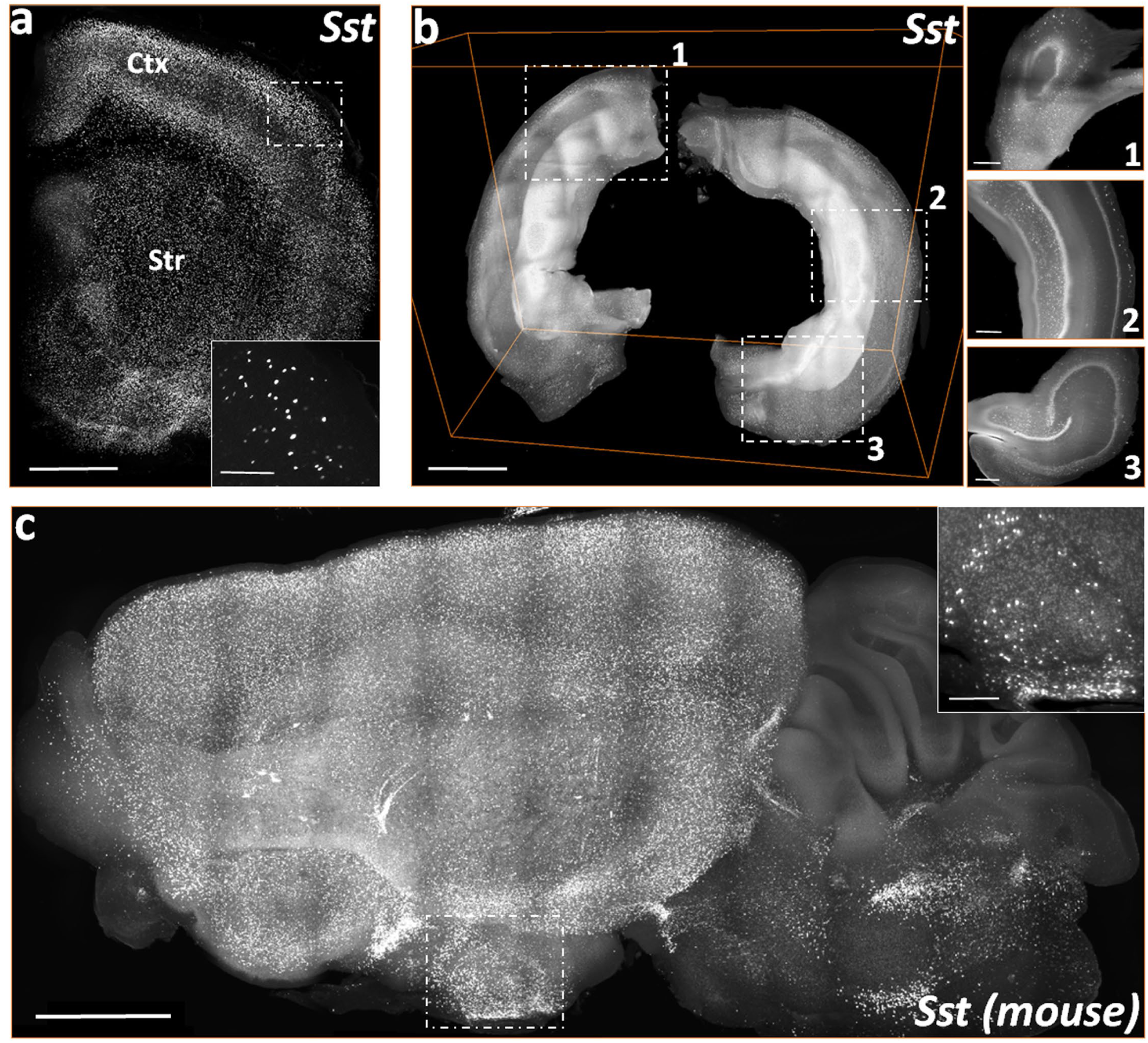
3D visualizations of COLM acquired stacks of S*st* mRNA labelling in the rodent brain cleared using iDISCO^+^. (a) A representative stack of rat forebrain hemi-slice corresponding to a 2 mm thick region between bregma = 2.00 mm and 0.00 mm, rostro-caudally. (b) Volume rendering of an intact rat hippocampus. Inset, *xy*-plane views of 1) dorsal, 2) medio-lateral, and 3) ventral hippocampus. (c) 3D volume rendering of a mouse left hemisphere (∼4 mm thick, sagittal view). Scale bars (µm) – (a) 300 (Inset-100); (b-c) 250 (Inset-100); (d) 500 (Inset-200); (e) 1000 (Inset-300); (f) 1000 (Inset-200).

A COLM-acquired volume of a rat forebrain (Figs. 4a, supplemental movie ESM_6) and brainstem hemi-slices (Fig. 4b, supplemental movie ESM_7) show the expression pattern of *Pvalb* mRNA. The mRNA expression pattern of *Th* in the ventral tegmental and substantia nigra regions of the rat brain slice can be seen in the COLM-acquired volume (Fig. 4c, supplemental movie ESM_8). The confocal microscope-acquired hemisphere volume in Fig. 4d, supplemental movie ESM_9 show the *Dbh* mRNA expression in the caudal half volume of the rat locus coeruleus (LC) consisting of the posterior pole and the middle part spanning the dorso-ventral axis. Following the successful validation in fresh-frozen rodent brain samples, we tested HCR FISH in iDISCO^+^-processed postmortem human brain tissues (∼1mm thick). The HCR-based labelling failed for the *PVALB* and *CALB* mRNAs in the temporal cortex slices. However, the cerebellum slices showed adequate signal for both of the above transcripts. *CALB* mRNA labeling is observed in most of the Purkinje cells (Fig. 4e, supplemental movie ESM_10) while the *PVALB* mRNA labelling is seen in the Purkinje cells as well as in the basket and stellate cells of the molecular layer (Fig. 4f, supplemental movie ESM_11).

**Fig. 4.**
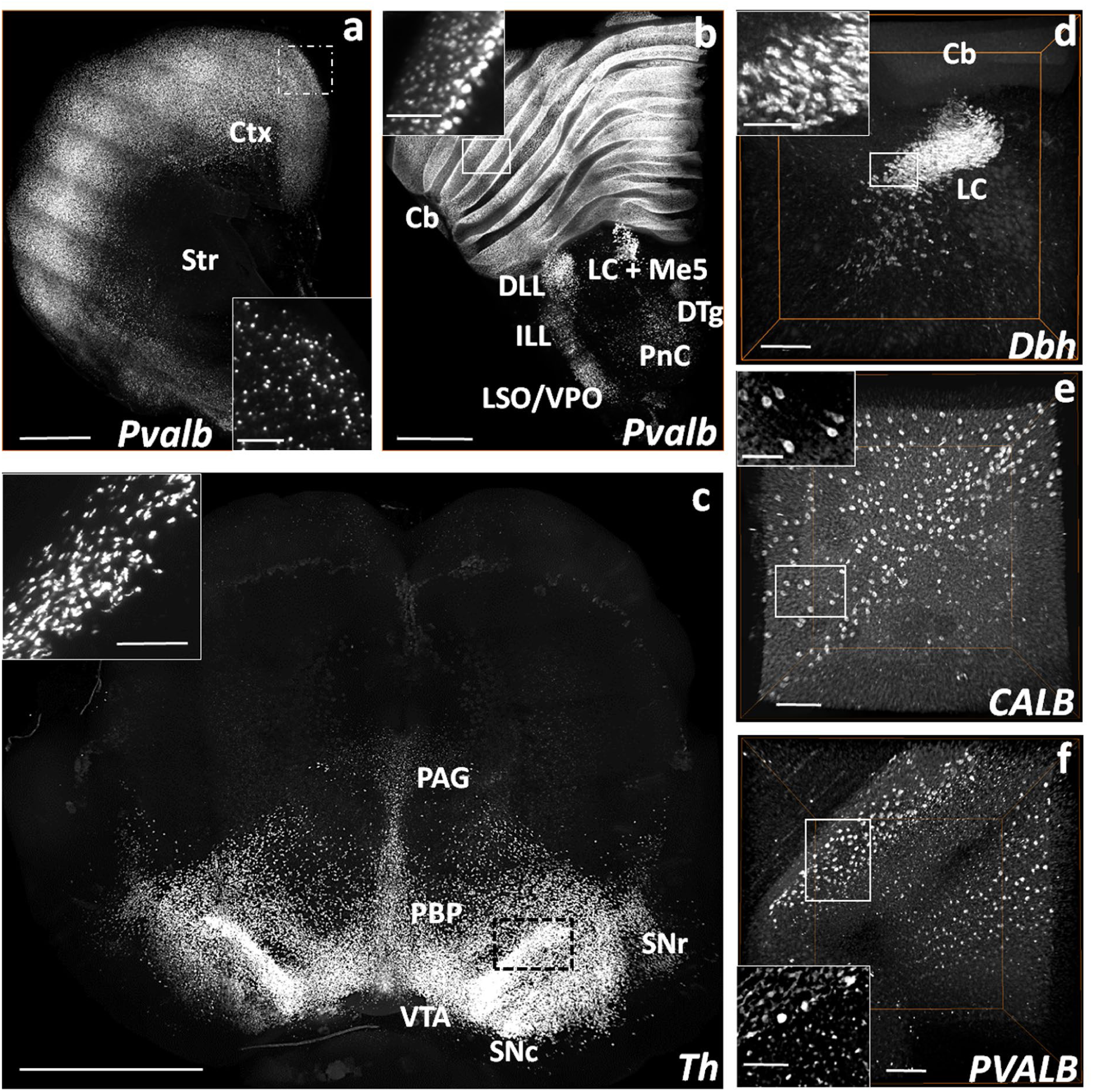
Volume rendered visualizations of *Pvalb, Dbh*, and *Th* mRNA expressing neurons in the rat brain, and *CALB*^*+*^ and *PVALB*^*+*^ neurons in the postmortem human cerebellum following HCR FISH and iDISCO^+^. (a) COLM-acquired volume of *Pvalb* expressing neurons in a 2 mm thick rat cortex (Ctx) and striatum (Str) hemi-slice between bregma = 2.00 mm and 0.00 mm. (b) COLM-acquired image of *Pvalb* expression pattern in a 2 mm rat brainstem hemi-slice between bregma = −8.44 mm and −10.44 mm. (c) Confocal microscope-acquired image stack of *Dbh* expression in the locus coeruleus (LC) of rat brainstem hemi-slice between bregma = −9.72 mm and −10.32 mm. (d) COLM-acquired image of *Th* expression in a 2.5 mm thick rat mid-brain volume between bregma = −4.40 mm and −6.90 mm. (e) *CALB* and (f) *PVALB* expression in confocal microscope-acquired stacks of postmortem human cerebellum blocks (1500 µm thick). Cb-cerebellum, DTg-dorsal tegmental nucleus, ILL/DLL-intermediate/dorsal lateral lemniscus, LC+Me5-locus coeruleus + mesencephalic trigeminal nucleus, PnC-caudal pontine reticular nucleus, LSO/VPO-lateral/ventral superior olive, PAG-periaqueductal gray, PBP-parabrachial pigmented nucleus of the VTA, SNc/r-substantia nigra compacta/reticulata, VTA-ventral tegmental area. Scale bars (µm) - (a-b) 1000 (Inset-400); (c) 500 (Inset-100); (d) 1000 (Inset-200); (e-f) 500 (Inset-250).

### Multiplexing ability of HCR FISH in cleared iDISCO^+^ tissues

Next, we evaluated the ability of HCR FISH to detect multiple genes in cleared fresh-frozen samples. Our initial experiments using both 10 µm thin sections as well as CLARITY- and iDISCO^+^-processed thicker samples showed inconsistency in the fluorescence signal with the use of different fluorophore tagged hairpins. To determine whether these variations were fluorophore-based, we designed a set of probes with the same mRNA detecting sequence but with different initiators, e.g., probe1-B1 and probe1-B2 (Choi et al. 2010). Using multiple initiator-specific DNA hairpins tagged with different fluorophores (e.g., B1H1-AF647:B1H2-AF647; B2H1-AF488:B2H2-AF488), we were able to detect and directly compare the HCR signal of the same gene in different laser channels. Fig. 5 shows confocal microscope-acquired HCR FISH images of *Dbh* mRNA in the LC region using hairpins conjugated with AF-488, AF-594, and AF-647 in 20 µm thin sections. Hairpins tagged with AF-647 produced a high S/N ratio (Fig. 5d), whereas hairpins tagged with AF-488 or AF-594 showed a low S/N ratio (Fig. 5b and 5c). A COLM-acquired image of iDISCO^+^-processed thicker tissues showed similar differences in fluorescence signal (Fig. 6a *vs.* 6b; 6c vs. 6d). Altogether these indicate a possible difference in the stability of fluorophores AF-488 and AF-594 or their conjugation stability with DNA hairpins, in comparison to the AF-647. We also tested the efficiency of DNA hairpins conjugated with alternative fluorophores FITC and Cy3 from IDT. However, both Cy3 conjugation (Fig. 6d *vs.* 6e) and FITC-conjugated hairpins (data not shown) failed to produce a consistent and comparable S/N ratio compared to that of AF-647 in the iDISCO^+^ samples.

**Fig. 5.**
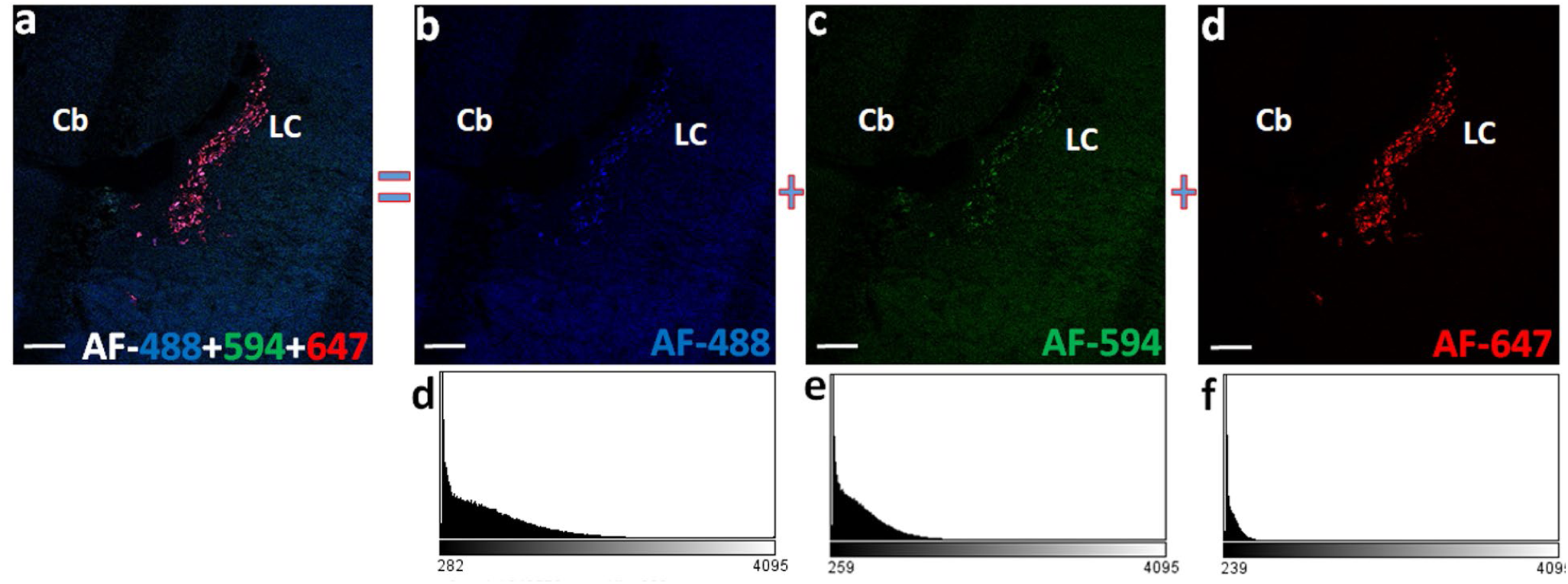
Fluorophore based differences in the fluorescence output imaged using confocal microscope. (a) *Dbh* expression in the rat locus coeruleus (LC) following multiplexed HCR FISH using a set of common mRNA binding probes and 3 different hairpins AF-488, AF-594 and AF-647. Relatively weak fluorescence and high background recorded with the use of AF-488 (b) and AF-594 (c) tagged DNA hairpins in comparison to the strong fluorescence with low background following the use of hairpins conjugated with AF-647 (d). Screenshots of the histogram (d, e, and f) for their respective (b, c, and d) images, generated on the Olympus FluoViewer software depicts the low S/N ratio with the use of AF-448 and AF-594 compared to a high S/N ratio with the use of AF-647. Images are pseudocolored for the overlaying purpose. Scale bars-200 µm.

**Fig. 6.**
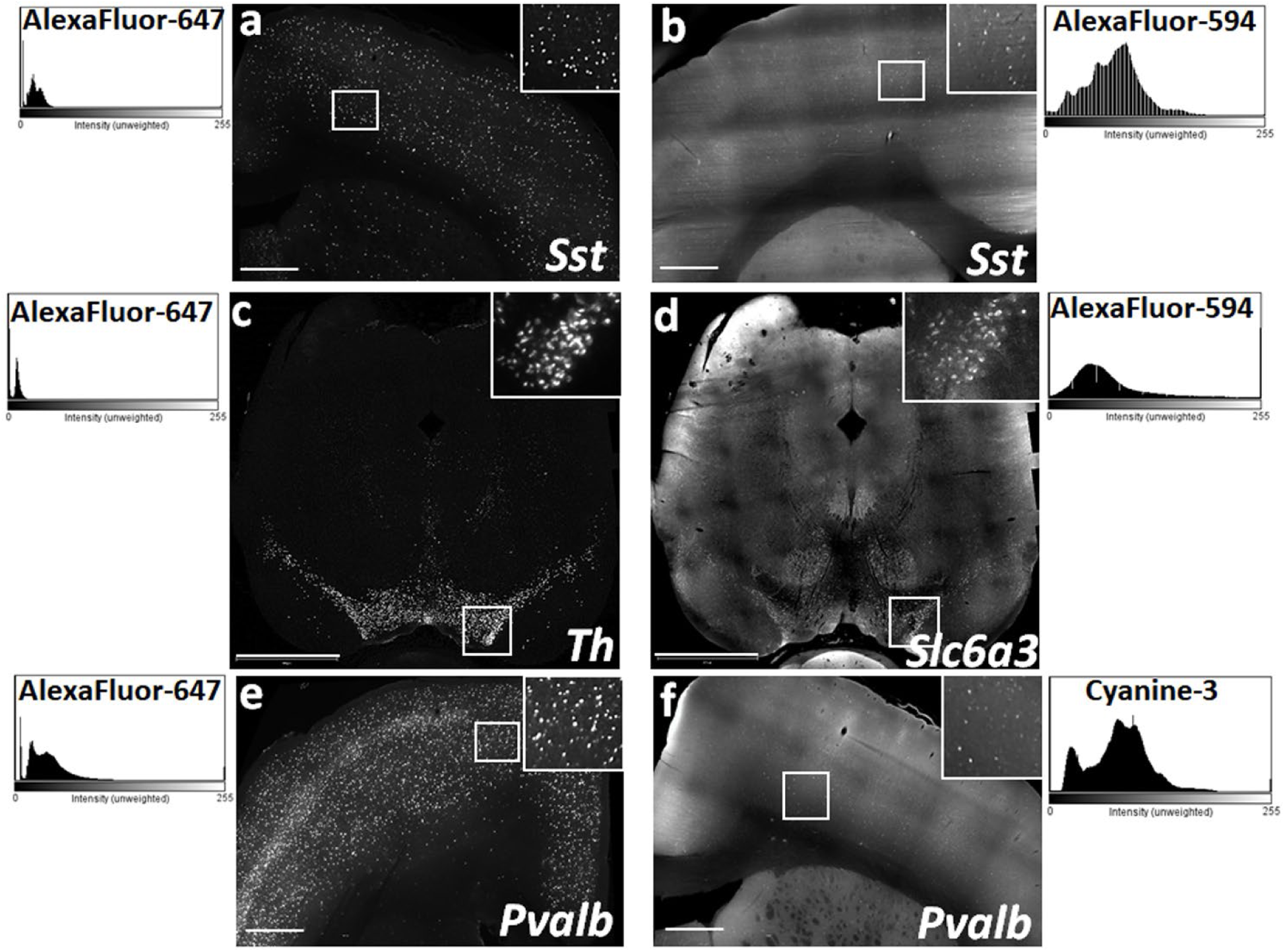
Fluorophore based differences in the quality of fluorescence output imaged using light sheet microscope. Representative *xy*-plane images of *Sst* labeled neurons in rat cortical slices using DNA hairpins conjugated with (a) AF-647 and (b) AF-594. Representative *xy*-plane images of rat mid-brain slices following multiplexed HCR FISH for genes with identical expression levels-(c) *Th*, detected with AF-647 and (d) *Slc6a3* (DAT) detected with AF-594. Comparison of the *Pvalb* signal in rat cortical tissue using DNA hairpins conjugated with (e) AF-647 and (f) Cy3. ImageJ generated histograms on the respective top corners, indicate high S/N ratio with the use of AF-647 in comparison to the low S/N ratio with AF-594 or Cy3. Scale bars-(a, b, e, f) 500 µm; (c-d) 1200 µm.

### Application of volumetric HCR FISH to assess the effect of GR overexpression on basal levels of *Sst* mRNA in mouse cortex

To establish a quantitative application of HCR FISH in combination with clearing methods, we compared the cortical expression of *Sst* transcripts between WT (n=5) and GRov (n=5) mice. Estimation of *Sst***^+^** neurons per mm^3^ (neuronal volume density) showed a significantly lower number of detected neurons in the cortex of GRov vs. WT mice in both CLARITY (two-tailed t(7) = 2.35, p = 0.051; Fig. 7a *vs.* 7b; 7e *vs.* 7f; 7i), and iDISCO^+^ samples (two-tailed t(10) = 2.84, p = 0.017; Fig. 7c *vs.* 7d; 7g *vs.* 7h; 7i). Normalized mean intensity of the detected *Sst*^+^ neurons was significantly lower in GRov cortex comparison to WT in CLARITY samples (two-tailed t(7) = 4.82, p = 0.002) but not in iDISCO^+^-processed samples (two-tailed t(3.5) = 1.67, p = 0.179; Fig. 7j).

**Fig. 7.**
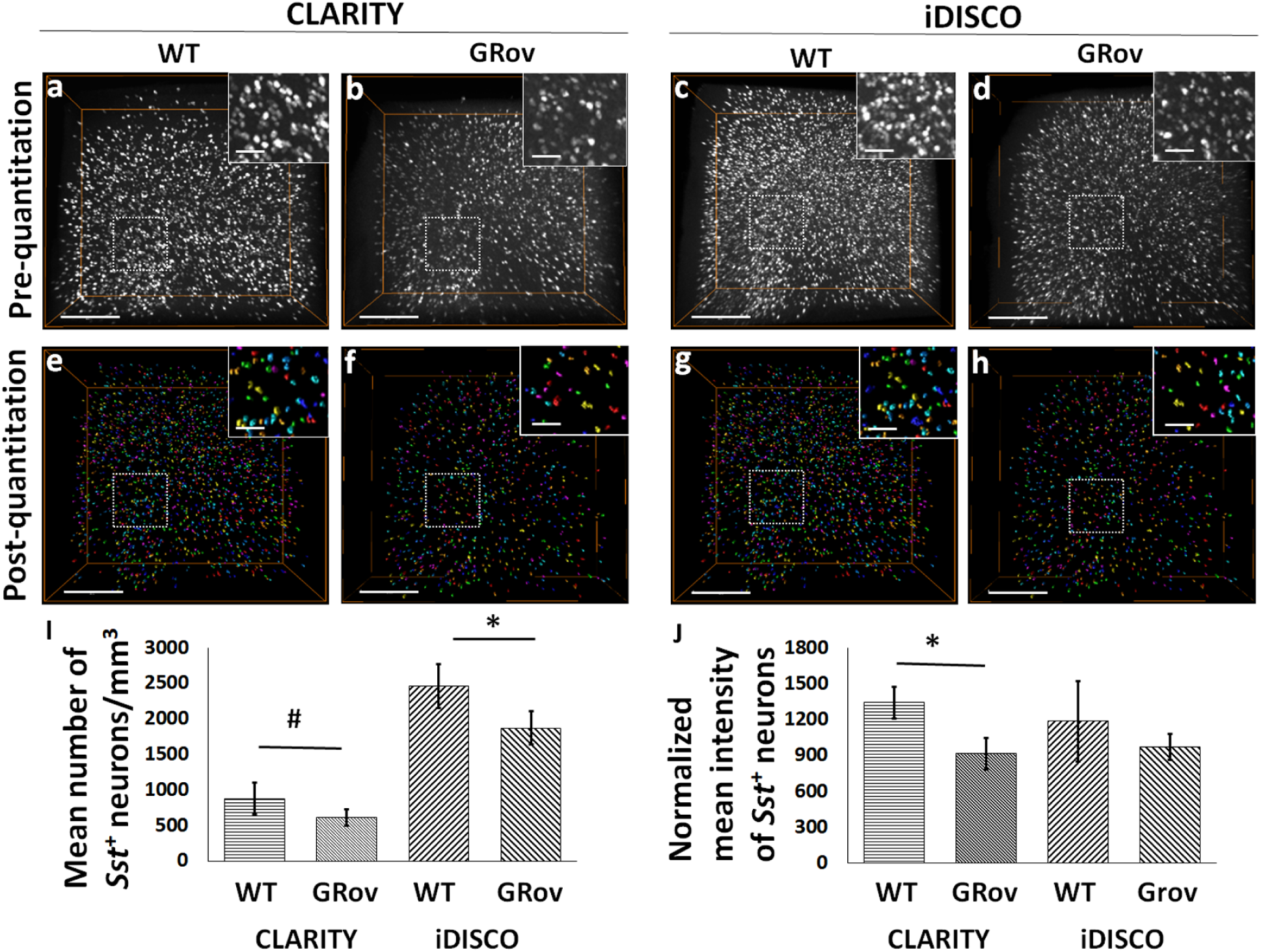
GR overexpression negatively affects the expression of *Sst* in the analyzed cortical ROI of mice. 3D volume-rendered confocal images show qualitative visualization of *Sst*^+^ neuronal density in CLARITY-(a) WT, (b) GRov; and iDISCO^+^-(c) WT, (d) GRov. Surface-rendered images show post-quantitation volume density of *Sst*^+^ neurons in CLARITY-(e) WT, (f) GRov; and iDISCO^+^-(g) WT, (h) GRov. (i) Bar-diagram shows the significant difference in the number of *Sst*^+^ neurons between WT and GRov samples processed through CLARITY (cWT *vs.* cGRov) or iDISCO^+^ (iWT *vs.* iGRov). (j) Bar-diagram shows significant difference in the normalized mean intensity of *Sst*^+^ neurons between cWT and cGRov. #p=0.051; *p<0.05. Scale bars-300 µm (Inset bars-100 µm).

## Discussion

In the current study, we sought to assess the compatibility and efficiency of HCR FISH in fresh-frozen brain samples cleared using different approaches and select the suitable platform to map and analyze transcripts. Based on the factors like clearing capability, target biomolecule fixation, ability to perform ex-vivo labeling, fluorescence retention and imaging depth, we explored only two transparency techniques, CLARITY (Chung et al. 2013;) with modifications (Yang et al. 2014; Tomer et al. 2014; Krolewski et al. 2018), and iDISCO^+^ (Renier et al. 2014). We chose fresh-frozen rodent brains as starting material to optimize steps like RNA fixation, probe permeability, hybridization and tissue clearing conditions in intact samples. Fresh-frozen tissues can be used to study a wide array of biomolecules (e.g., transcripts, proteins, metabolites) using methods like *in situ* hybridization, immunohistochemistry, PCR, western blotting, spectroscopy etc. Traditionally, fresh-frozen tissues have been preferentially used as starting material for both radioactive- and digoxigenin-based *in situ* hybridization. On the other hand, fixed tissues from live animal perfusions have superior retention of selective biomolecules especially proteins, and allow better lipid removal leading to efficient sample transparency. However, fixation limits the utilization of tissues for numerous biochemical techniques. Altogether, flash freezing of fresh brain samples provides the most flexible utilization of the tissue and it is certainly the most ethical method employed for postmortem human brain collection for research purposes.

### Comparison of HCR FISH in fresh-frozen brain tissues cleared using CLARITY or iDISCO^+^

The time and efficiency of sample clearing using CLARITY was dependent upon nature of tissue (rodent *vs.* human) and sample thickness, whereas, iDISCO^+^ processing yielded optimal transparency within a standard time period (Renier et al. 2014) in the fresh-frozen brain samples irrespective of tissue origin or size. Typically, under passive clearing conditions in 4% SDS (Tomer et al. 2014), lipid removal time for a typical 2-4 mm thick slice of 2% acrylamide + 4% PFA perfused rodent brain ranges from 1-2 weeks. In our experience, it could take up to 2-3 weeks to achieve a comparable transparency in fresh-frozen immersion-fixed rodent slices. Additional clearing time is required for postmortem human brain tissues of similar thickness mainly due to the higher tissue density and extensive myelination. In terms of imaging depth capability, we consistently achieved depth of 1500-2500 µm in iDISCO^+^ samples whereas in most of the CLARITY samples, mRNA labelling could be recorded up to only 700-1000 µm deep.

Quantitatively, both CLARITY- and iDISCO^+^-processed mouse cortical samples showed a comparable mean signal intensity (*Sst* mRNA expression), suggesting a similar efficacy for the combination of HCR FISH and fresh-frozen immersion-fixed samples of a given thickness. However, the mean tissue background intensity (with or without including the *Sst* signal) was significantly lower in CLARITY compared to iDISCO^+^ samples. The final steps of the iDISCO^+^ method including dehydration with alcohol, de-lipidation with DCM, and RI matching with DBE, modify the tissue as evident from shrinkage and hardening. These modifications cause local variations in the RI in cleared iDISCO^+^ samples depending upon the regional content and morphology (e.g., ventricles, edges, and fiber dense regions) and contribute to higher tissue background compared to CLARITY samples.

CLARITY processing induced expansion causes the fluorescence signal from *Sst*^*+*^ neurons to be recorded on relatively more number of pixels which, in turn, allows the detected neurons to be represented by a wide range of pixel intensity, making more accurate intensity-dependent measurements possible. In contrast, the tissue shrinkage with iDISCO^+^ processing allows detection of more neurons per unit volume and representation of *Sst*^*+*^ neurons by fewer pixels. Majority of these pixels have mean intensity equal or higher than the threshold value used as cutoff. Nevertheless, this iDISCO^+^ limitation is offset by time-saving and more efficient computational resource usage to acquire relatively larger samples and perform volume analysis.

Analysis of the HCR FISH signal as a function of image z-depth reveals a significant attenuation of the signal in CLARITY samples. Mean pixel number quantitation shows that neurons located at deep layers consist of significantly fewer pixels than those located towards top layers. In contrast, fresh-frozen samples with similar thickness when processed with iDISCO^+^ showed relatively uniform labelling of *Sst*^+^ neurons across the z-depth. A combination of sample transparency and tissue size change leads to the difference in the z-depth signal pattern between these two clearing procedures. Considering the tissue shrinkage, better transparency, and resulting ability to image deeper, the iDISCO^+^ method allowed the acquisition of larger tissue volumes compared to CLARITY.

### Validation of HCR FISH in iDISCO^+^-cleared rodent and postmortem human brain

We successfully validated our optimized combination of HCR FISH and iDISCO^+^ methods by imaging the expression of a few abundant to moderately expressed genes (e.g., *Sst, Pvalb, Th, Dbh*) in intact fresh-frozen rodent brain tissues. However, HCR FISH failed to detect low mRNA copy number genes e.g., adrenergic receptor alpha-1A and 1B or genes expressed in sparse neuronal population e.g., phenylethanolamine-N-methyltransferase. This indicated a low sensitivity of HCR FISH towards less expressing genes especially in the large tissues subjected to clearing procedure. By increasing the number of probe-pairs to hybridize a large portion of mRNA length could further amplify the fluorescence signal and better mRNA preservation one can further amplify the signal from less expressing genes.

Next, we investigated the potential of HCR FISH to label transcripts, *SST, VIP, PVALB*, and *CALB* in the postmortem human brain tissues. We could not detect any adequate signal in the cortical slices irrespective of the gene or clearing method used. In contrast, HCR FISH in iDISCO^+^ cleared human cerebellum blocks produced fluorescence signal from *PVALB* and *CALB* transcript labelling. Differences in the regional tissue density and mRNA expression levels are likely reasons for the observed differences between cortex and cerebellum samples. Insufficient clearing of tissues with high density (extensive myelination) could reduce the probe permeability and optical transparency as we observed with postmortem human cortical slices which took longer to clear with CLARITY method than cerebellum blocks. Low expression or preservation of mRNA in the tissue could dictate the sensitivity of HCR FISH. The presence of lipofuscin-induced autofluorescence across a wide spectral range further affects the imaging ability, especially from less abundant genes. We tried commonly used reagents such as cupric chloride, sodium borohydrate, sudan black and trueblack™ (Biotium, Inc., Fremont, CA, USA) to quench the autofluorescence in combination with clearing methods, but did not succeed. This necessitates further optimization of HCR FISH and clearing methods to capture low quality and lower copy number of transcripts in the postmortem human brain, along with the lipofuscin-induced autofluorescence.

### Fluorophore based differences in the HCR FISH signal output

We found that the multiplexing ability of HCR FISH is critically dependent upon the fluorophores which produce different quality of the fluorescence output. For example, AF-647 tagged hairpins produced the most consistent fluorescence signal and highest S/N ratio and proved to be the most resistant to photobleaching under extended imaging sessions using both confocal and light sheet microscopes. In contrast, hairpins tagged with AF-488 and AF-594 failed to produce as consistent and comparable S/N ratio across the experiments. One possible reason for the differences in the fluorescent signal output could be the relative instability of AF-488 and AF-594 molecules themselves or their conjugation to the DNA hairpin when compared to that of AF-647. According to the proprietary manufacturer Thermofisher Scientific, the AF-647 fluorophore has superior photostability in buffers and mounting media than both AF-488 and AF-594, suggesting an inherent difference in the chemistry. Based on our experience, these differences are more evident when HCR FISH is performed in combination with tissue clearing methods that involves longer incubations in high salt concentration buffer (5x SSCTw) and tissue clearing solvents like methanol, DCM or DBE, as opposed to thin section experiments. It is also possible that the higher quantum efficiency of the sCMOS cameras (COLM) or photomultiplier tubes (confocal microscope) in the spectral region of 450 to 600 nm contributes to the observed noise as seen with AF-488 or AF-594. Further experiments are required to establish the exact cause of such fluorophore-based differences in signal output as there appear to be several possibilities. We also explored the efficiency of DNA hairpins conjugated with alternative fluorophores FITC and Cy3, but these failed to produce a comparable signal to that of AF-647. We compared the HCR FISH signal using AF-647-conjugated hairpins purchased from either IDT or Molecular Instruments, Inc. to test whether vendor differences in conjugation chemistry in generating DNA hairpins makes a difference. We observed less consistency in the fluorescence signal across the FISH experiments using IDT-synthesized AF-647-conjugated hairpins (data not shown), as compared to hairpins from Molecular Instruments, indicating that different conjugation chemistry may be important. Moreover, when immunochemistry was performed using secondary antibodies tagged with the same fluorophores in combination with tissue clearing methods, the above mentioned differences in fluorescence output were not observed (Krolewski et al. 2018). This further indicates that conjugation chemistry likely plays a significant role in the stability of the fluorophores tagged on to the DNA hairpins vs. antibodies.

### Forebrain glucocorticoid receptor overexpression reduces the basal level of cortical *Sst* mRNA expression

Finally, we used this optimized volumetric quantitative HCR FISH to compare the basal expression of *Sst* transcripts in the cortex of WT and GRov mice. Our lab has previously shown that transient early-life, as well as constitutive, GR overexpression in forebrain causes profound long-term alterations in the region-specific transcriptome and emotional reactivity (Wei et al. 2004; Wei et al. 2007; Hebda-Bauer et al. 2010; Wei et al. 2012). GRov mice exhibit significantly higher levels of total GR mRNA and approximately 78% more GR protein in the forebrain than wild type (WT) controls (Wei et al. 2004). The brains used in the present study were from mice that had GR overexpressed in forebrain from early life throughout adulthood (from when the Ca2+/calmodulin-dependent protein kinase promoter becomes active around the 2nd week of life until the animals were sacrificed). Our findings showed that GR overexpression beginning in early life negatively affects the basal expression of *Sst* expression in the adult cortex, as shown by a significantly lower *Sst*^+^ neuronal number density when compared to the WT samples, processed either through CLARITY or iDISCO^+^. A similar trend in the normalized mean intensity suggests low *Sst* transcript copy number in GRov cortices compared to WT. Our observation is consistent with previous studies showing that GR overexpression may amplify the overall glucocorticoid response in the cortex which, in turn, negatively affects the *Sst* expression in the rodent cortex (Papachristou et al. 1994). Nevertheless, this basal level effect of forebrain GR overexpression on cortical *Sst* expression should be validated with an increased sample size and additional quantitative methods.

## Conclusion

The HCR FISH is compatible with both CLARITY and iDISCO^+^ clearing methods for the detection of abundant to moderately expressed mRNAs. The iDISCO^+^-based tissue clearing approach provided the more suitable and versatile platform for analyzing transcripts in fresh-frozen and immersion-fixed tissues. Nevertheless, iDISCO^+^-mediated tissue modifications, namely shrinkage and hardening, induce a morphology dependent variable RI which tends to increase the background, especially if the probe and hairpin concentrations are not well titrated for larger samples. Tissue shrinkage with iDISCO^+^ helps the probe and hairpin to penetrate relatively large tissue volumes in less time, and enables more efficient image data processing and management. On the other hand, CLARITY images were better resolved with higher S/N ratios than iDISCO^+^ images likely due to tissue expansion and improved probe and hairpin penetration. The superior resolution of CLARITY images also helps minimize the variations in intensity-based quantitative comparisons. However, less efficient clearing and restricted imaging depth in fresh-frozen samples limit the use of CLARITY for thin or small tissues. HCR FISH in cleared samples provide a unique opportunity to study postmortem human brains, which are a valuable resource for understanding the underlying mechanisms of psychiatric disorders. Nonetheless, postmortem human brain presents unique challenges (e.g., tissue and RNA quality as a function of postmortem interval, higher tissue density mainly due to extensive myelination and presence of lipofuscin-induced autofluorescence) which demands further optimization of HCR FISH and clearing methods. Overall, we conclude that the benefits of iDISCO^+^ make it more suitable for HCR FISH based transcript analysis in intact rodent and postmortem human brain samples with further optimizations needed to enhance the sensitivity of HCR FISH.

## Supporting information

Supplemental movie ESM_1

Supplemental movie ESM_2

Supplemental movie ESM_3

Supplemental movie ESM_4

Supplemental movie ESM_5

Supplemental movie ESM_6

Supplemental movie ESM_7

Supplemental movie ESM_8

Supplemental movie ESM_9

Supplemental movie ESM_10

Supplemental movie ESM_11

## Acknowledgements

This work was supported by the Pritzker Neuropsychiatric Research Consortium, the Hope for Depression Research Foundation, National Institute of Health R01MH104261, Office of Naval Research Grant N00014-12-1-0366 and National Institute on Drug Abuse U01DA043098. The authors have no conflicts of interest to declare. The authors would like to thank Ms. Jennifer Fitzpatrick, Mr. Evan Hughes, Ms. Claire Barcelo, and Mr. Hui Li for their technical assistance.

## Compliance with Ethical Standards

### Ethical approval

“All animal care and procedures performed in studies involving animals were in accordance with the ethical standards of the guide for the Care and Use of Laboratory Animals: Eighth Edition (revised in 2011, published by the National Academy of Sciences), and approved by the University of Michigan Committee on the Use and Care of Animals.

“This article does not contain any studies with human participants performed by any of the authors.”

### Informed consent

Postmortem human brains were collected by the Brain Donor Program at the University of California, Irvine with the consent of the relatives of the deceased.

## Supplemental material legends

**Supplemental movie ESM_1**. 3D visualization of differences in the HCR-FISH labeled *Sst* signal between CLARITY (left panel) and iDISCO^+^ (right panel) processed mouse cortical slices. First half of this movie shows a series of *xy*-plane images moving in *z*-direction of confocal acquired representative stacks, followed by maximum intensity projection (mip) 3D pre-quantitation volume and post-quantitation surface rendering of *Sst* neurons. Second half of the movie shows a series of *xz*-plane images moving in *y*-direction, followed by mip 3D pre-quantitation volume and post-quantitation surface renderings of *Sst* neurons. This animation highlights the higher number density of detected *Sst* neurons and uniform labeling across the z-plane in iDISCO^+^ compared to CLARITY.

**Supplemental movie ESM_2**. 3D visualization of HCR FISH-labeled *Sst* expressing neurons in a rat forebrain hemi-slice cleared using iDISCO^+^. This movie first shows a series of xy-plane images acquired on a COLM followed by a mip 3D volume rendering. The characteristic layer specific expression pattern of *Sst* in rat cortex can be observed in this movie.

**Supplemental movie ESM_3**. Volume rendering of a rat hippocampus showing S*st* expression pattern in the iDISCO^+^ cleared intact tissue and labelled using HCR FISH method. This movie shows a series of xy-plane images acquired on a COLM followed by 3D mip volume rendering.

**Supplemental movie ESM_4**. 3D visualization of a mouse left hemisphere showing *Sst* expressing neurons across the entire rostro-caudal extent of the brain. Post HCR FISH labelling, brain tissue was cleared using the iDISCO^+^ method and imaging was performed on a COLM.

**Supplemental movie ESM_5**. *Sst* expression in an iDISCO^+^ cleared sample of a mouse left hemi-brainstem. COLM-acquired images are first rendered as a series of xy-planes progressing in the medio-lateral direction and subsequently as mip 3D volume rendering.

**Supplemental movie ESM_6**. Volume rendering of HCR FISH-labeled *Pvalb* expressing neurons in a rat forebrain hemi-slice. This representative COLM-acquired image stack of an iDISCO^+^-cleared brain shows the predominant cortical expression of *Pvalb*.

**Supplemental movie ESM_7**. 3D visualization of HCR FISH-labeled *Pvalb* expressing neurons in a rat brainstem hemi-slice showing the expression pattern in cerebellum and several nuclei. iDISCO^+^ cleared and COLM-acquired image stacks are first projected as a series of xy-planes moving in a rostral-caudal-rostral direction, then as a mip 3D volume rendering.

**Supplemental movie ESM_8**. 3D visualization of an iDISCO^+^ cleared rat mid-brain slice showing the expression of *Th* mRNA in the SNr/c, VTA, PBP and PAG regions. This movie first shows the rostro-caudal progression of a series of *xy*-plane images acquired on the COLM followed by a mip 3D volume rendering.

**Supplemental movie ESM_9**. *Dbh* expressing neurons in the rat locus coeruleus are shown here first as a series of *xy*-plane images acquired on a confocal microscope followed by a mip 3D volume. This representative brainstem hemi-slice was processed through HCR FISH and cleared using the iDISCO^+^ method.

**Supplemental movie ESM_10**. 3D visualization of HCR FISH-labeled *CALB* expressing neurons in the postmortem human cerebellum block. Confocal microscope-acquired image stacks are rendered first as a series of *xy*-plane images followed by a mip 3D volume rendering.

**Supplemental movie ESM_11**. *PVALB* expressing neurons in the postmortem human cerebellum block are visualized in this movie as a series of *xy*-plane images then by a mip 3D volume. This representative block was cleared using iDISCO^+^ and images were acquired on a confocal microscope.

